# Single-cell RNA-seq analysis profiling characterizes differences in cell composition and physiology between normal tissue, treatment naive, and cisplatin-treated ovarian cancer

**DOI:** 10.1101/2024.01.02.573967

**Authors:** Fang Guo, Zhi Yang, Jalid Sehouli, Andreas M. Kaufmann

## Abstract

**Background:** Intense efforts have focused on identifying heterogeneity of the cellular composition in ovarian cancer. However, tissue composition and physiological conditions of cancer cells in cisplatin-sensitive ovarian cancer remains largely unknown. Moreover, comparisons of different cellular states in normal tissue, in treatment naive ovarian cancer, and in cisplatin-treated tissue after adjuvant therapy of cisplatin-sensitive ovarian cancer at the single-cell level might offer clues for ovarian cancer treatment and prevention of cisplatin-resistance formation.

**Methods:** Single-cell transcriptome sequencing of a cisplatin-treated ovarian cancer was performed. Data sets of non-tumorous ovarian tissues and treatment-naive ovarian cancer were downloaded from the European Genome-phenome Archive (accession number EGAS00001004987). Quality control, batch effect correction, integration, and clustering analysis of the integrated single-cell transcriptome data was done. Cell subsets were annotated based on surface marker phenotypes. Finally, the proportions of subclusters, the immune cell population, and the potential biological processes among different cellular states were compared.

**Results:** Sixteen distinct cell subsets were identified from the integrated single-cell transcriptome sequencing data of a pool of all tissues. The composition of the three different tissue types was characterized. The proportion of fibroblasts in cisplatin-treated ovarian tumor was remarkably lower than in treatment-naive ovarian tumor (1.33% vs. 13.53%, p < 0.05). Moreover, each subject’s sample had differing relative proportions of the identified cell types. In primary untreated ovarian cancer, the prevalent immune cells were B cells and myeloid-related immunosuppressive M2 macrophages. However, there were less B cells and myeloid-related immunosuppressive M2 macrophages after cisplatin-treatment, while significantly more T cells were found. The physiological cellular state in primary untreated ovarian tumors was associated with dysfunctional gene expression and modulation of cellular homeostasis, while cells from cisplatin-treated tumor showed more activation of immune and inflammatory genes as compared to healthy ovarian tissue.

**Conclusion:** Our molecular gene expression analysis allowed for the separation and identification of differences in normal ovarian tissues, treatment-naive, and cisplatin-sensitive ovarian cancer cell populations at single-cell resolution. We identified different cell type composition and discriminative marker expression concerning specific cell subsets and identified differences among their physiological cell states. This knowledge may open new possibilities for elucidating important pathogenetic features and therapeutic strategies for treating ovarian cancer.

## Introduction

Ovarian cancer, one of the most fatal and aggressive cancer types, have increased in recent years (Siegel, Miller et al. 2019). The treatment is primarily cytoreductive surgery followed by adjuvant chemotherapy. Platinum-based chemotherapy is the current standard treatment for ovarian cancer (Kelland 2007). It is effective in high-grade serous ovarian cancer (HGSOC), and around 70% of HGSOC respond to a platinum-based chemotherapy. Platinum sensitivity (or lack thereof) is a major determinant of prognosis. the 10-year survival rate was improved by more than 10% by the introduction of platinum-based combination therapy. (Kelland 2007). However, 50% of these patients will develop recurrence (Damia and Broggini 2019) due to development of platinum resistance. Intense efforts have focused on identifying various cell types in platinum-resistant cancers. However, defining the cellular composition and cellular processes of platinum-sensitive cancers remains largely unknown. Moreover, a better understanding of the differences between treatment-naive versus cisplatin-treated cellular composition and physiology in platinum-sensitive ovarian cancers can be informative about sensitivity mechanisms. Studying these differences will allow to elucidate better platinum-sensitive ovarian cancer pathogenesis, treatment responses, and characterize the endogenous heterogeneity of cells in platinum-sensitive ovarian cancers.

In addition, HGSOC often develop fatal recurrence due to development of resistant cells that grow out after treatment. Enhanced DNA repair, and tumor microenvironment alterations like hypoxia, acidosis, physical blockade, and reactions of cancer stem cells and immune cells are factors that contribute to the effect of chemotherapy and the clinical outcome of ovarian cancer treatment. Several initiatives, such as the Australian Ovarian Cancer Study (AOCS) (Patch, Christie et al. 2015) and The Cancer Genome Atlas (TCGA)(Network and Bell 2011), have identified four molecular subtypes of the HGSOC by applying conventional bulk gene expression analysis: the mesenchymal, immunoreactive, differentiated, and proliferative HGSOCs. These findings highlight the importance of the tumor microenvironment and cellular composition in HGSOC and point out the necessity to explore its heterogeneity more accurately which may elucidate factors for disease progression and therapy responsiveness or resistance.

To study various cell types and the composition among cell populations, single-cell RNA sequencing (scRNA-seq) was performed in this study. One recent scRNA-seq research studied the ovarian tumor heterogeneity at cellular resolution (Shih, Menzin et al. 2018). Another study using single-cell RNA sequencing technology identified eight diverse clusters and showed the similarity between embryos and tumors in some clusters based on their gene expression (Zhao, Gao et al. 2021). These novel studies provided insight and enhanced our understanding of ovarian carcinogenesis. However, to understand the prevention modalities, progression under therapy, and development of more effective treatments, platinum-sensitive ovarian cancers’ cell types and functions must first be well understood.

ScRNA-seq enables to explore the transcriptomic diversity of tumors and to better understand the most relevant chemotherapy-induced processes at an unprecedented level in various cell types. Data from scRNA-seq also addresses the interplay between cancer cells and the tumor stroma microenvironment. This study compared the characteristics and differences among non-tumorous ovarian tissue, treatment-naive ovarian cancer, and cisplatin-treated ovarian cancer showing treatment sensitivity at single-cell resolution. Comparing treatment-naive ovarian cancer and cisplatin-treated ovarian cancer cell populations may help to understand how tumor cells respond to the stress of chemotherapy, the relation to the tumor microenvironment, and the significance of genomic instability and transcriptomic adaptation. Here we characterize and compare the cellular composition and the physiologic state in a cisplatin-treated therapy-responsive tumor with untreated ovarian cancer and with normal adjacent non-tumorous ovarian tissue. Indeed, a better understanding of this knowledge is likely to be central for improving therapeutic outcomes.

## Materials and Methods

### Ovarian cancer tissue collection

The high-grade serous ovarian cancer (HGSOC) tissue was obtained from a 46 year old multiparous woman with regular menstrual cycles. The imaging suggested that she got primary ovarian cancer. The patient underwent chemotherapy before surgery. Written informed consent was obtained from the patient, and this project received ethical approval from the ethical committee of Baoan Maternal and Children Health Hospital (LLSC 2020-03-28).

### Tissue digestion and cell preparation for single-cell RNA-sequencing

The ovarian cancer tissues were incubated in DMEM high glucose culture medium supplemented with 3 mg/ml collagenase D (Worthington-biochem, Lakewood, NJ, USA) at 37°C for 45 min. Then the tissues were digested in 1 mg/ml collagenase D in DMEM high glucose culture medium at 37℃ for three hs. After that, the cell suspension was transferred to a 40 µm cell filter and washed with 0.4% BSA in PBS. Further pipetting was done to fully disperse cell aggregates into single cells.

### Single-cell RNA-seq library construction and sequencing

GemCodeTM Single Cell platform (10X Genomics, Pleasanton, CA, USA) was used to determine the transcriptomes of single cells. Briefly, single-cell suspensions were brought to a concentration of 1000 cells/µl, and approximately 10000 cells were loaded, followed by GEM-RT reaction and cDNA amplification. Single-cell library preparation was performed by attaching P7 and P5 primer sites in Illumina bridge amplification of the cDNA, following the manufacturer’s instructions. Finally, the Illumina HiSeq4000 was used to sequence the library into 150-bp paired-end reads.

### Analysis and visualization of scRNA-seq data

Quality control was done using the Cell ranger 2.0.1 and the results were aligned to the hg38 human genome assembly. Poor quality cells were excluded. Cells that expressed fewer than 100 genes or more than 4000 genes were excluded, and cells with a mitochondrial gene percentage over 15% were discarded. Then, we performed an initial normalization per cell basis using the Seurat package, followed by log transformation and scaling. Data obtained from different platforms were uniformed to to exclude the quality factors. Through the function “FindAllMarkers” or “FindMarkers” in Seurat packages, DEGs were identified. As marker genes, significant DEGs were selected from genes with an average expression difference > 0.5 natural logs (P < 0.05). Enriched gene oncologies (GO) terms of marker genes were identified. The non-tumorous ovarian tissues and treatment-naive ovarian cancer data sets were downloaded from the European Genome-phenome Archive under accession number EGAS00001004987.

## Results

### ScRNA-seq reveals sixteen distinct cell populations in the three different tissues

For chemotherapy the patient received platinum (500 mg) combined with liposomal paclitaxel (240 mg) intravenously every three weeks. The serum CA125 levels were detected before and after chemotherapy. Cytoreductive surgery was performed after three cycles of chemotherapy. A sustained decrease of ≥50% in the serum CA125 levels was observed. Thus, the patient’s cancer was defined as platinum sensitive.

To investigate differences among healthy non-tumorous ovarian tissues, treatment-naive ovarian cancer, and cisplatin-sensitive ovarian cancer, we re-analysed the treatment-naive ovarian cancer and corresponding adjacent healthy ovarian tissue data from the European Genome-phenome Archive under accession number EGAS00001004987. These data contained a total of fresh biopsies from 7 treatment-naive patients during primary cytoreductive surgery or diagnostic laparoscopy, including primary treatment-naive ovarian tumor and the normal adjacent tissue. Our cell quality result was satisfactory for performing single-cell sequencing. The Uniform Manifold Approximation and Projection (UMAP) was performed, resulting in a total of sixteen putative cell clusters by using classic markers and differentially expressed genes (DEGs) (Figure 1-3). The sixteen cell clusters are designated as follows: (0) ovarian stroma cells, (1) epithelial cancer cells, (2) T cells, (3) myeloid cells, (4) muscle cells, (5) fibroblasts, (6) endothelial cells, (7) epithelial cancer cells, (8) fibroblasts, (9) epithelial cancer cells, (10) B cells, (11) myeloid cells, (12) myeloid cells, (13) fibroblasts, (14) epithelial cancer cells, (15) myeloid-related fibroblasts. Same cell types occur reputedly due to differences in their expression profiles. Representative markers for these cell clusters are revealed in Figure 2, representing combinations of genes to distinguish different cell populations. Overall, these results identified and quantitated various cell types of platinum-sensitive ovarian tumor treatment-naive ovarian tumors, and the normal adjacent tissues.

### Characterization of the composition of the three different tissues

Each cell type had successfully been annotated with known marker gene expression. Moreover, the relative abundance of potential cell types in the healthy non-tumorous ovarian tissue, the treatment-naive ovarian tumor, and the cisplatin-treated ovarian tumor, were compared. Our results showed that the proportions of cell types were quite different. As shown in Figure 4, in a healthy state, ovarian stroma cells (37%), muscle cells (19%), myeloid cells (11%), and fibroblasts (10%) were the major cell types present. Compared with healthy ovarian tissue, the relatively higher abundance in the treatment-naive ovarian tumor were in epithelial cancer cells (30%), ovarian stroma cells (19%), and myeloid cells (11%). Overall, healthy ovarian tissue was enriched for tissue stroma, while tumors were enriched for epithelial cell types. In cisplatin-treated ovarian cancer, ovarian stroma cells (44%) and T cells (33%) were the major cell types, indicating that the immune response may play a role in chemotherapy-sensitive cancer treatment.

Another interesting result was that the proportion of endothelial cells between treatment-naive ovarian tumor and the cisplatin-treated ovarian tumor was similar (3.82% v.s. 4.18%, p > 0.05), while the proportion of fibroblasts in the cisplatin-treated ovarian tumor was remarkably lower than that in treatment-naive ovarian tumor (1.33% vs. 13.53%, p <0.05). These results indicated that reduction of cancer-associated fibroblasts during chemotherapy may be associated with sensitivity to treatment with cisplatin.

### Differences in immune cell populations

All cells were grouped in 16 clusters representing seven major cell types based on marker gene expression: muscle cells, epithelial cancer cells, fibroblasts, T cells, myeloid cells, B cells, and tumor cells (Figure 5, Table 1). Our results showed that, among the healthy non-tumorous ovarian tissues, treatment-naive ovarian cancer, and cisplatin-treated ovarian cancer, each subject had relative proportions of different cell types. In ovarian cancer, the proportion of the epithelial cancer cells was 41.4%, however, after chemotherapy, the proportion of the epithelial cancer cells in cisplatin-treated ovarian cancer was only 4.88%, indicating that this tumor sample was indeed platinum-sensitive as the epithelial cancer cells were eliminated by platinum. These results were consistent with the serum CA125 levels which were decreased after cisplatin-treatment. Interestingly, the immune cells were quite different in these samples. In untreated ovarian cancer, the primary immune cells were B cells (2.38%). After chemotherapy, the major immune cells were T cells (33.0%), and the relative proportion of B cells was reduced to only 1.04%.

### Identification of potential biological process differences in immune cells

We sought to determine further the functional characteristics of the primary immune cell types. We had already found that the immune cell population might play a vital role during the response to chemotherapy. All immune cell types were merged to perform a more detailed comparison. In immune cells, ten new subclusters were identified, and gene ontology (GO) analyses were performed to explore potential biological and physiological processes.

These ten immune cell subclusters were grouped as: B cells, CD4/CD8 T cells, NK cells, Macrophages/DCs, plasmablast, and pDCs (Plasmacytoid dendritic cells) (Figure 6). B cells were present in both untreated tumor and cisplatin-treated ovarian cancer, and there were almost no B cells in the healthy ovarian tissue. There were CD4/CD8 T cells and NK cells present in all three different tissues, albeit in differing abundance. The CD4/CD8 T cells were a major component in cisplatin-treated ovarian cancer, while NK cells were mainly in untreated tumor. Our results show that the most different characteristics between immune cells in untreated tumor and the healthy ovarian tissue or the cisplatin-treated cellular composition are the macrophages/DCs cluster. There are more macrophages/DCs in tumor tissues compared with the healthy and the cisplatin-treated samples. The cells in cluster 1 are the majority from the treatment-naive tumor cells, and the cells in cluster 2 are the majority from the cisplatin-treated cells. Here, we sought to further determine functional characteristics in these populations. Differentially expressed genes (DEGs) in the cluster 1 population were enriched in GO terms such as “myeloid cell activation”, “neutrophil activation”, “myeloid leukocyte mediated immunity”, and “antigen processing and presentation,” suggesting their involvement in the regulation of immune suppression in the tumor immune microenvironment.

DEGs in the cluster 2 population were enriched in GO terms such as “immune response”, “inflammatory response”, and “immune effector process”, suggesting their involvement in the cellular cytotoxic immunity and inflammatory processes (Figure 7). These results indicated that this tumor state is related to the dysfunction of modulation of cellular homeostasis, and cisplatin-treated cells are related to activation of immune and inflammatory genes.

## Discussion

The major obstacle in ovarian cancer therapy is its intrinsic or quickly developing resistance against cisplatin treatment. Therefore, much effort is focused on investigation of mechanisms of cisplatin resistance in ovarian cancers. The scRNA-seq analysis of ovarian cancer cells in cisplatin-sensitive ovarian cancers has not been employed to our knowledge. Further, a detailed understanding of the cellular composition and its alteration during therapy response in healthy ovarian tissue and in a therapy-induced cell states in cisplatin-treated ovarian cancer at the single-cell level is critical to gain further insight into ovarian cancer pathogenesis and treatment options. We are the first to report the differences between healthy ovarian tissue, treatment-naïve pimary ovarian cancer, and chemotherapy-induced cellular tumor composition and cellular physiology based on scRNA-seq analysis. We investigated these three tissue conditions and their cellular states at single-cell resolution. Importantly, we characterized the changes between healthy tissue, the untreated malignant cancer and the short-term cisplatin-treated tumor to elucidate the pathogenesis, changes by transformation, and alterations by treatment, in order to contribute to improved treatment of human ovarian cancer.

In recent years, researches have shown that the tumor microenvironment plays a crucial role in tumor immune suppression, distant metastasis, and the targeted therapy response (Paluskievicz, Cao et al. 2019, Schulz, Salamero-Boix et al. 2019). The tumor microenvironment is a highly complicated system composed of tumor cells, cancer-associated stromal cells, fibroblasts, endothelial cells, and diverse immune cells like macrophages, dendritic cells and tumor infiltrating lymphocytes. All these cells secrete metabolites, cytokines, immunosuppressive factors, along with multiple other signaling molecules. To investigate the microenvironment in different tissues, proportions of cell types were compared among normal ovarian tissue, primary treatment-naive ovarian tumor, and cisplatin-treated ovarian cancer tissue. Our results show that normal ovarian tissue was enriched for stroma cells, muscle cells, and fibroblasts according to the expected tissue architecture that has limited numbers of epithelial cells. The treatment-naive ovarian tumor contained mainly epithelial cancer cells, stroma cells, and myeloid cells. In contrast, T cells were primarily found in the cisplatin-treated ovarian cancer tissue sample. One interesting finding was that the proportion of fibroblasts in cisplatin-treated ovarian cancer tissue was remarkably reduced compared to treatment-naive ovarian tumor. Fibroblasts, which may interact with tumor cells, are the essential components of the tumor microenvironment. Thus, fibroblasts play critical roles in tumorigenesis and development, including angiogenesis, invasion, and metastasis (Bu, Baba et al. 2019, Liu, Han et al. 2019). Previous reports have shown that fibroblasts participate in multiple stages of tumor development through diverse signaling, including cytokines, chemokines, growth factors, and exosomes. Fibroblasts also induce immune evasion of cancer cells and facilitate tumor proliferation (Farhood, Najafi et al. 2019, Kobayashi, Enomoto et al. 2019). Our results are consistent with these previous reports. The proportion of fibroblasts in cisplatin-treated ovarian cancer tissue was remarkably reduced, indicating that the tumor microenvironment changes and in our case fibroblasts co-depletion with the epithelial cancer cells were correlated with sensitivity to treatment with cisplatin.

The tumor immune microenvironment has received increased attention related to the clinical prognosis of patients with tumors (Chen, Zhou et al. 2020). Distinct cell populations, including innate and adaptive immune cells, comprise most of the tumor immune microenvironment. Depending on the composition and activity of infiltrating immune cells, the tumor immune microenvironment also determines the state of the immune response (Zhang, Liu et al. 2020). Our results show that the immune cell composition and physiologic cellular states were quite different in the compared samples. In untreated primary ovarian tumor, the primary immune cells were B cells and NK cells, indicating that B cells and NK cells played essential roles in tumor immune suppression. There are conflicting roles for B cells in tumors in different researches. Some of the results showed that B cells in tumors could suppress antitumor immunity by producing IL-10, TGF-β, and IL-35, or by directly promoting tumor progression through the lymphotoxin/IKKα-BMI1 signaling pathway (Yuen et al., 2016). In contrast, some other studies have shown that B cells could improve antitumor effects by producing antibodies, secreting antitumor cytokines, and serving as antigen presenting cells (Yuen, Demissie et al. 2016). All these conflicting observations may be partially due to the heterogeneity of B cells in different tumor microenvironments. In our results, B cells are in high frequency in ovarian tumors representating a non-cytotoxic immunity or a TH2-biased immune response, while after chemotherapy they are depleted and the immune landscape swiched to a T cell-biased response.

Consistent with our finding of increased T cell frequenciess, others have also described the increased number of memory CD8+ T cells after chemotherapy (Liu, Tayob et al. 2022). Francesco F Fagnoni’s research (Fagnoni, Lozza et al. 2002) showed that the increase in T-cell proliferation was observed at the end of each chemotherapy cycle, and an effect of chemotherapy on B cells was not directly observed. Recent research also revealed an increased T cell response after administration of chemotherapy. Notably, the increase in T cell responses paralleled the decrease in CA125 levels which is consistent with an excellent response to chemotherapy. We speculate that these results potentially reflect the reinvigoration of suppressed effector and memory T cell populations with increased homeostatic proliferation.

Our results also revealed biological processes in tumor tissue and the cisplatin-treated ovarian cancer. The most pronounced GO terms of treatment-naive tumor cells were “myeloid cell activation”, “myeloid leukocyte-mediated immunity”, and “antigen processing and presentation”. Myeloid-related leukocytes have an immunosuppressive role in ovarian cancer which is associated with their ability to promote cancer growth, invasion, angiogenesis, immune evasion, and metastasis (Worzfeld, Pogge von Strandmann et al. 2017). However, B cells and myeloid-related leukocytes dropped after chemotherapy, while the T cell frequency increased. These results are consist with others’ results that the higher frequencies of myeloid-related leukocyte types from tumor-associated ascites strongly correlated with higher tumor grade (Yuan, Zhang et al. 2017). In our patient case, the ovarian cancer was cisplatin-sensitive, thus, the B cells and myeloid-related leukocytes decreased after chemotherapy. The GO terms of cisplatin-treated tumor cells were “immune response”, “inflammatory response”, and “immune effector process”. These results indicate that immune effector cells were activated and recruited into tumor tissues after chemotherapy. In previous studies it was found that fibroblasts can restrict the recruitment of immune effector cells into tumor tissues through the secretion of several chemokines (Ene–Obong, Clear et al. 2013). Moreover, the activated immune cells also secret some cytokines, such as interleukin (IL)-1β, and facilitate the recruitment of inhibitory immune cells and enhance immune suppression in the tumor immune microenvironment.

We acknowledge that the limited sample size of the available repository data sets and the single case of the cisplatin-treated patient imposes limitations on this study and the result interpretation. However, our results may provide clinically meaningful insights that may be relevant for immunotherapy of ovarian cancer. A better understanding of the immune cells infiltrating the tumor immune microenvironment will help to improve characterization of the treatment-naive and cisplatin-treated tumor tissue.

## Conclusion

In conclusion, single-cell RNA-seq analysis allowed for the separation and identification of the cellular populations in normal ovarian tissue, primary treatment-naive ovarian tumor, and of cisplatin-treated ovarian cancer tissue at single-cell resolution. Our research identified different cell types, their proportions in the cellular populations, and within the certain cell types different cellular physiological states. This methodology might offer clues for understanding ovarian cancer progression, help deepen our understanding of cancer immune suppressive and of immunotherapy-related mechanisms, and provide an insight into new therapeutic targets for treating ovarian cancer.

## Data availability statement

The datasets generated for this study are available on reasonable request to the corresponding author.

## Ethics statement

The studies involving human participants were reviewed and approved by the Ethics Committee of Baoan Maternal and Children Health Hospital. (LLSC 2020-03-28) The patient provided written informed consent before participating in this study.

## Author contributions

Conceptualization, F.G., Z.Y., B.W., J.S., A.M.K.; methodology, F.G.; software,B.W.; formal analysis, F.G., Z.Y., A.M.K.; resources, F.G.; data curation, F.G., Z.Y., B.W, A.M.K.; writing — original draft preparation, F.G.; Manuscript review and editing, A.M.K.; supervision, A.M.K.; Validation, F.G., Z.Y., B.W., A.M.K. All authors have read and agreed to the published version of the manuscript.

---

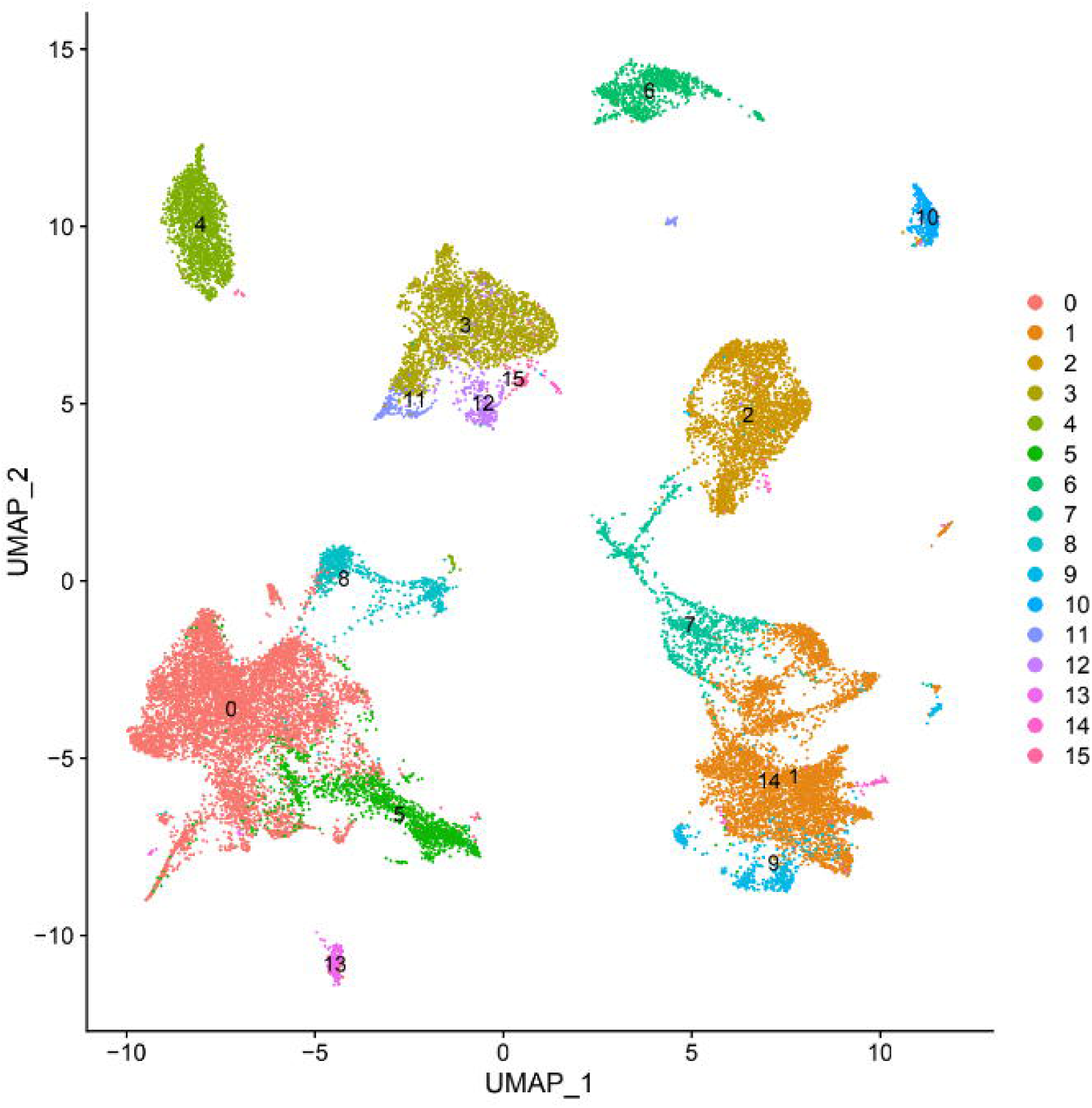

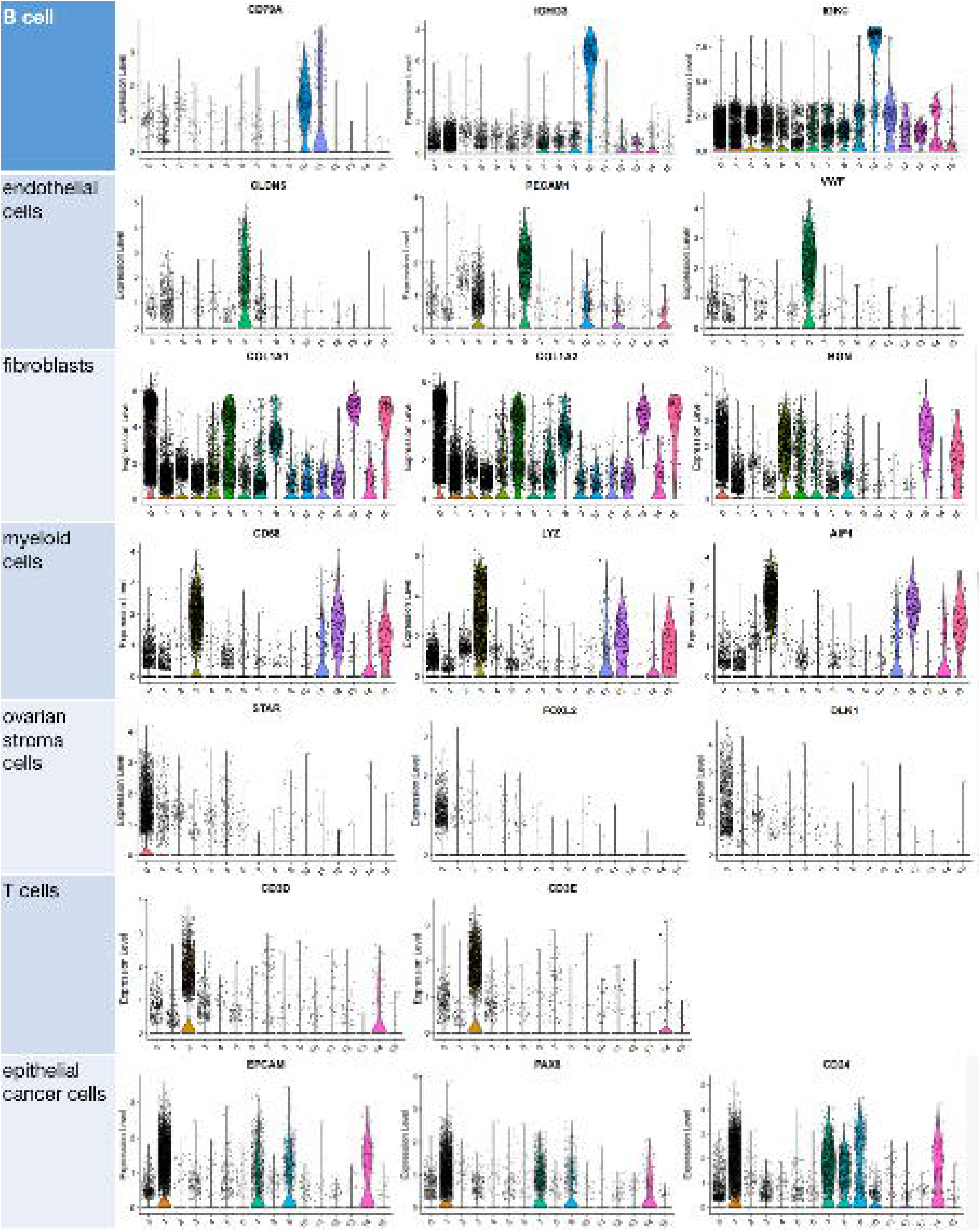

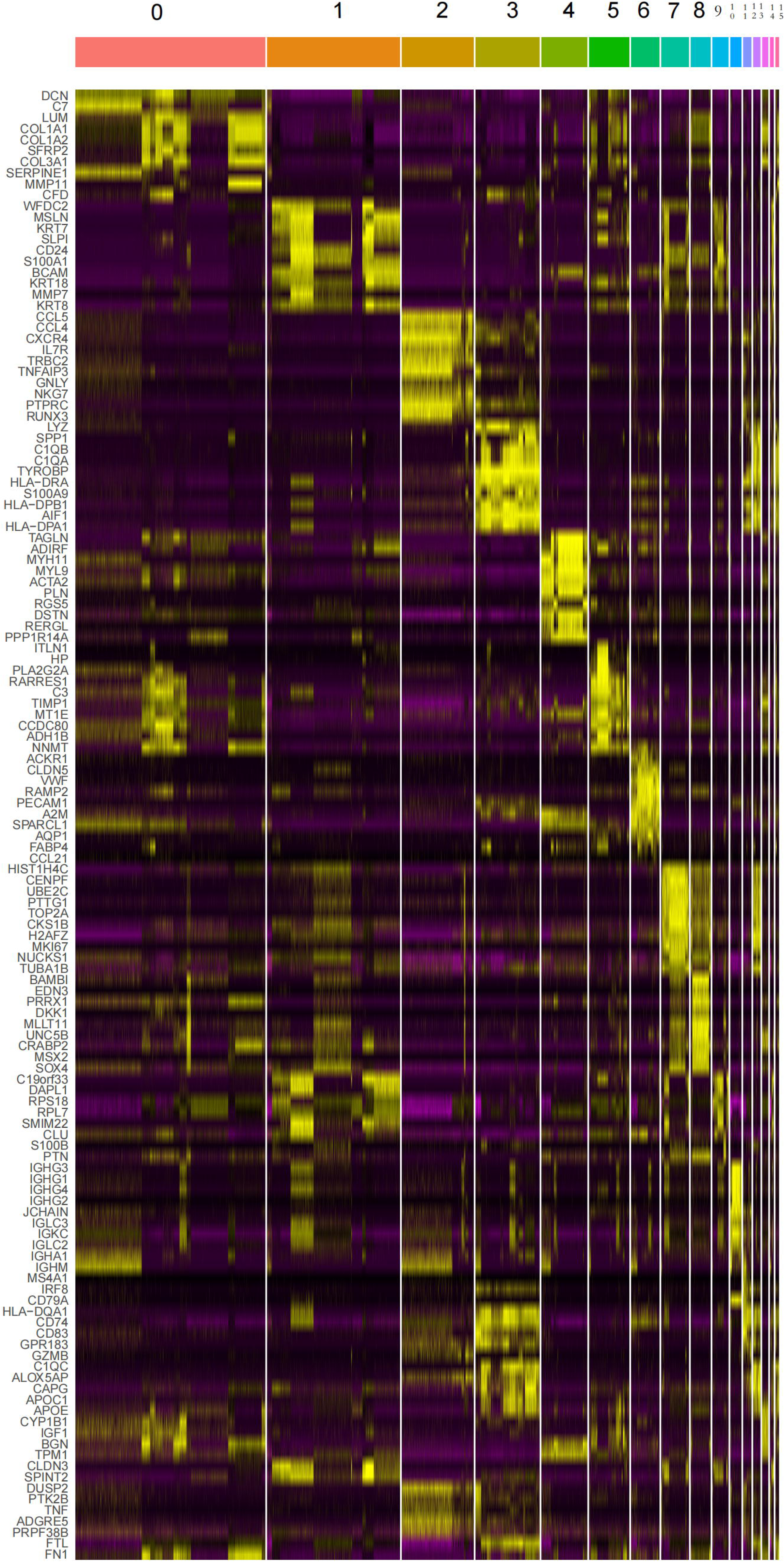

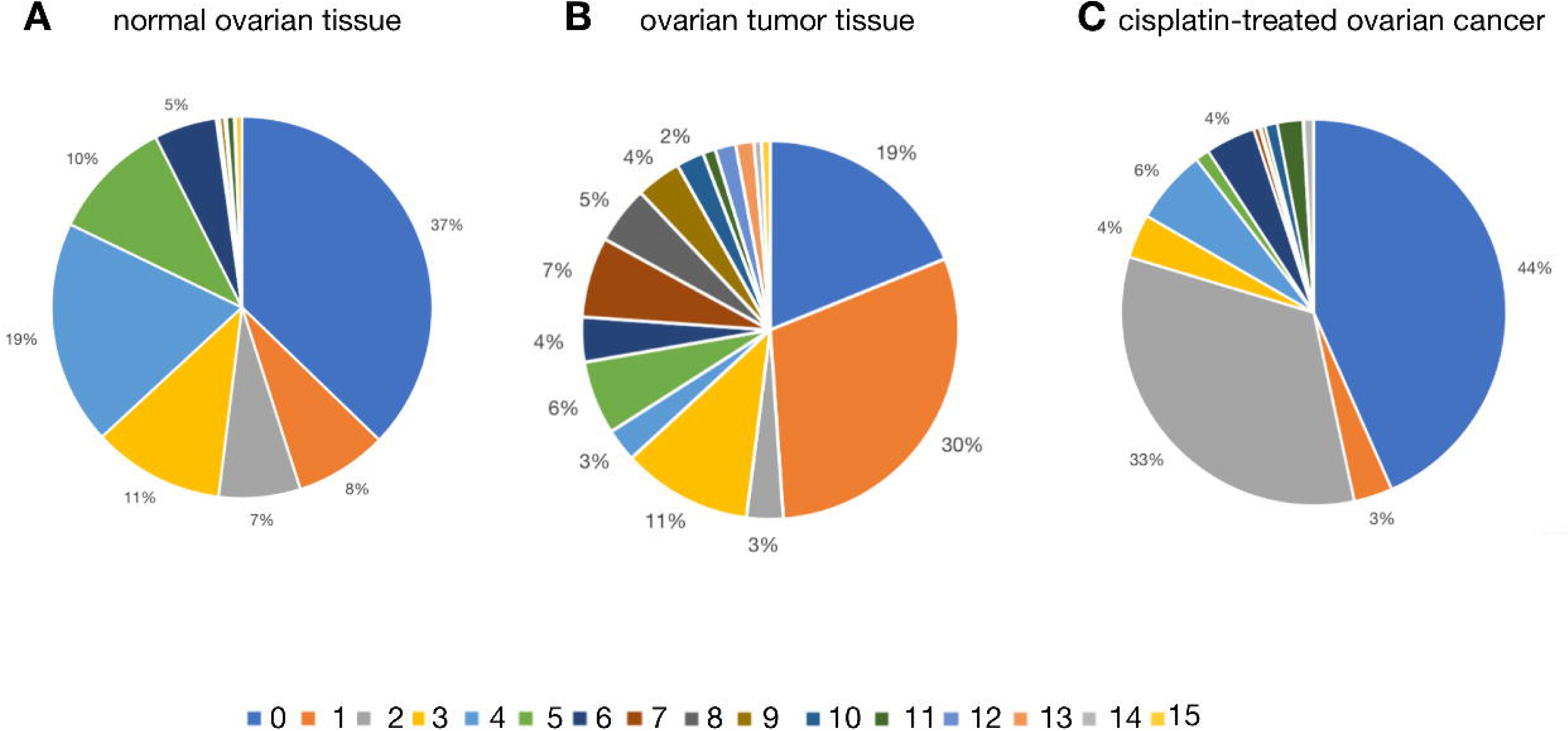

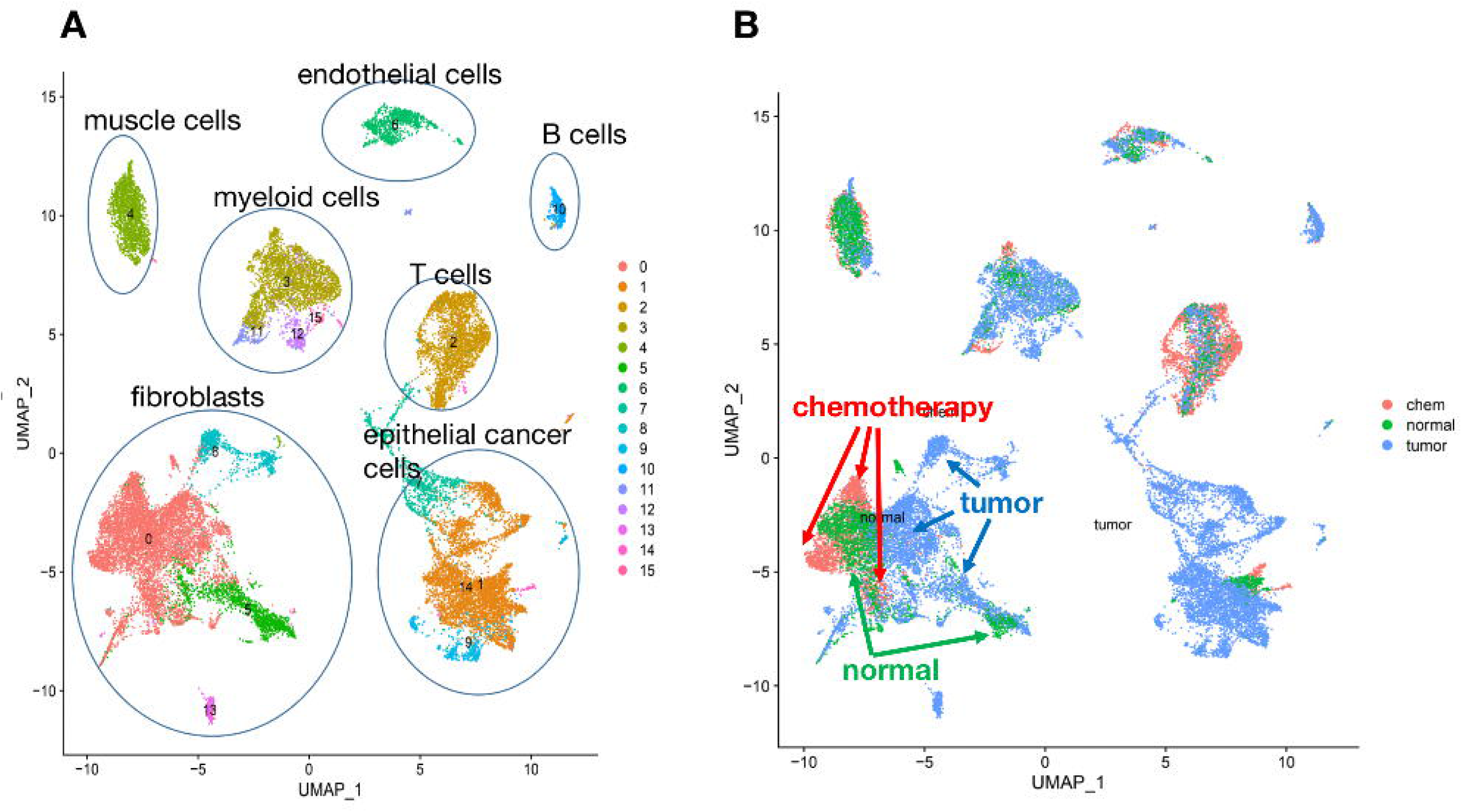

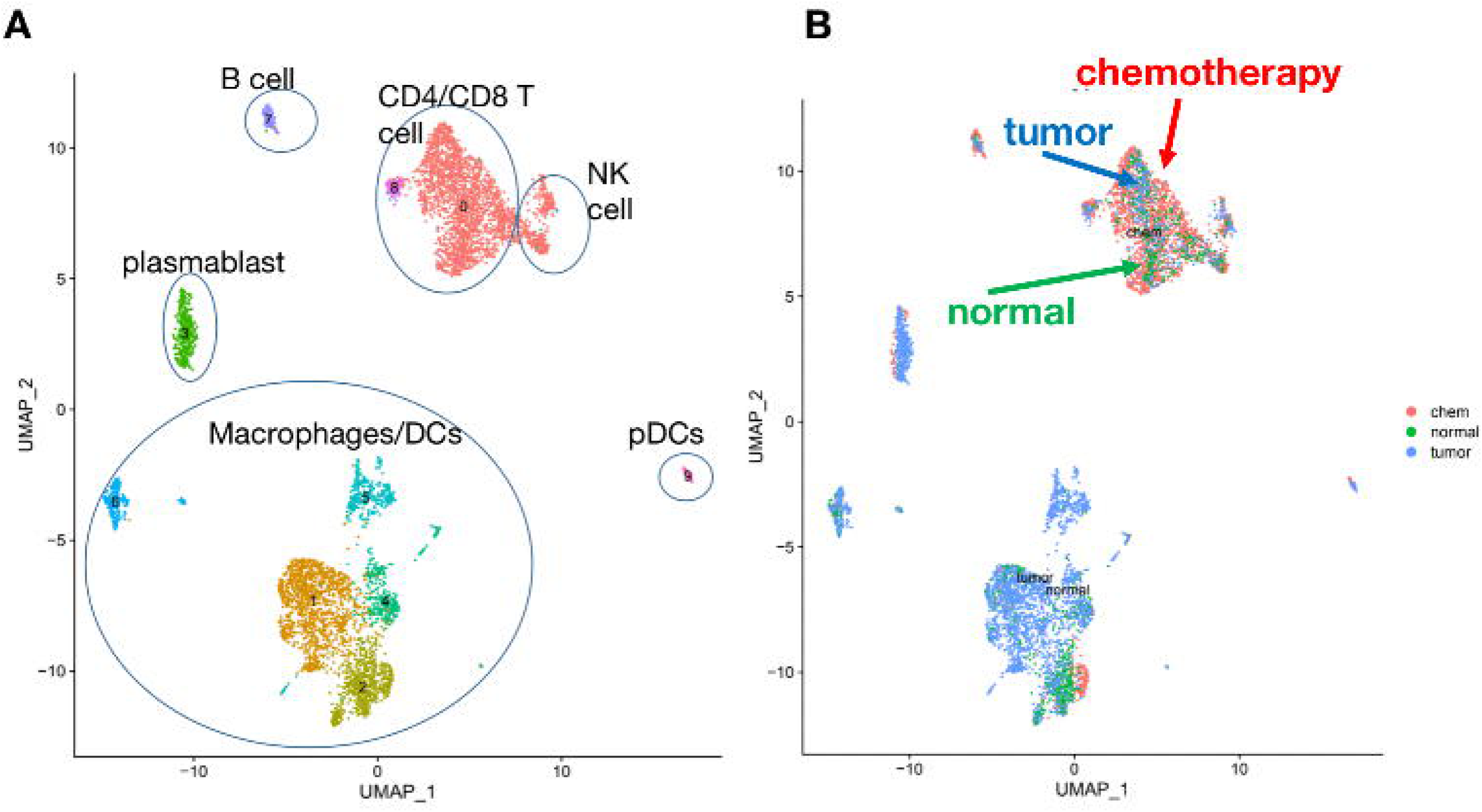

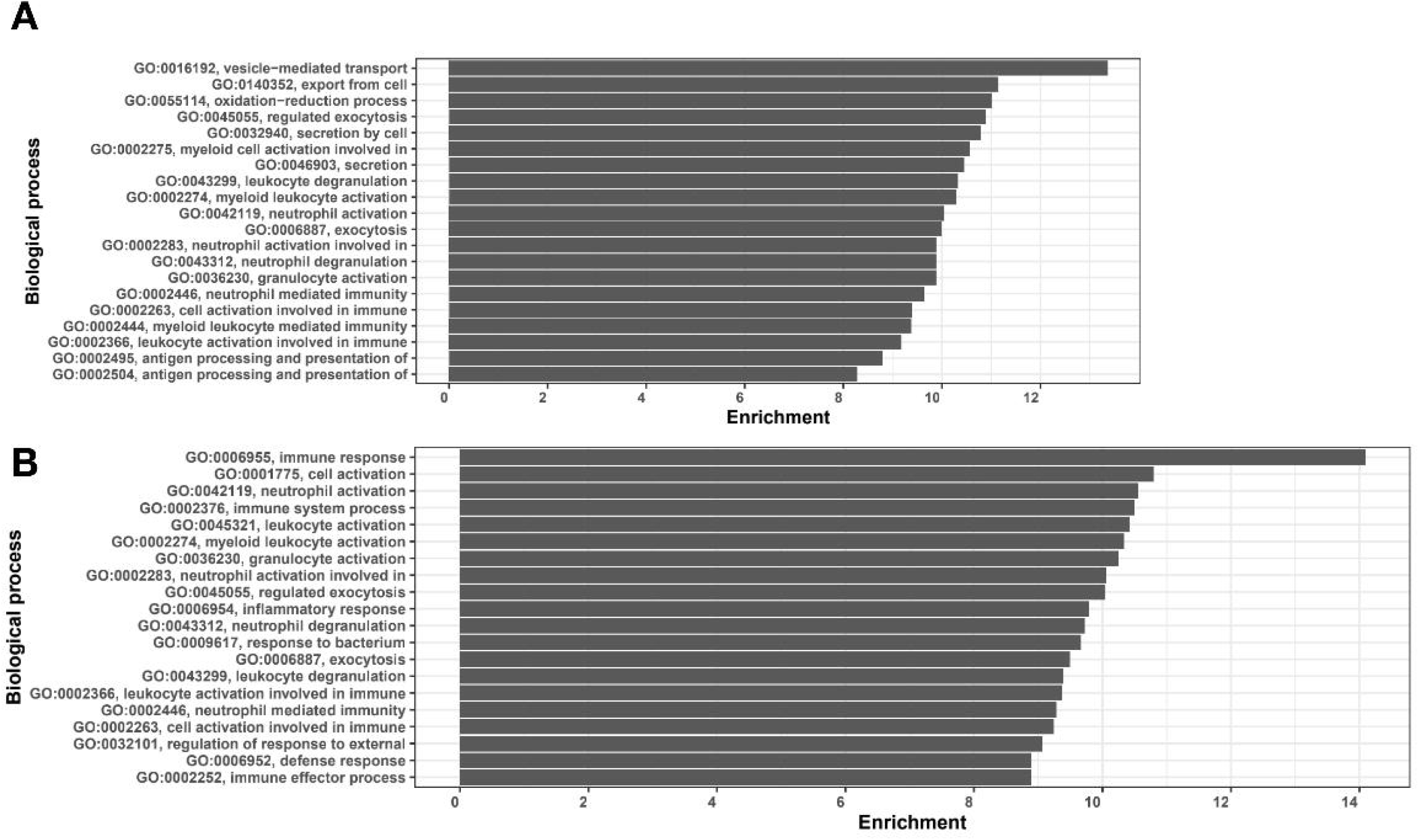

